# IL15/IL15Rα complex induces an anti-tumor immune response following radiation therapy only in the absence of Tregs and fails to induce expansion of progenitor TCF1+ CD8 T cells

**DOI:** 10.1101/2024.09.18.613691

**Authors:** Miles Piper, Jacob Gadwa, Chloe Hodgson, Michael Knitz, Elliott Yee, Yuwen Zhu, Keira Y. Larson, Christian Klein, Maria Amann, Anthony Saviola, Sana D Karam

## Abstract

**Background:** This work seeks to understand whether IL15-incorporating treatments improve response to radiotherapy and uncover mechanistic rationale for overcoming resistance to IL15 agonism using novel therapeutic combinations.

**Experimental Design:** Orthotopic tumor models of PDAC were used to determine response to treatment. IL15-/- and Rag1-/- mouse models were employed to determine dependence on IL15 and CTLs, respectively. Flow cytometry was used to assess immune cell frequency and activation state. Phospho-proteomic analyses were used to characterize intracellular signaling pathways.

**Results:** We show that the combination of radiation therapy (RT) and an IL15/IL15Ra fusion complex (denoted IL15c) fails to confer anti-tumor efficacy; however, a CD8-driven anti-tumor immune response is elicited with the concurrent administration of an aCD25 Treg-depleting antibody. Using IL15-/- and Rag1-/- mice, we demonstrate that response to RT + IL15c + aCD25 is dependent on both IL15 and CTLs. Furthermore, despite an equivalent survival benefit following treatment with RT + IL15c + aCD25 and combination RT + PD1-IL2v, a novel immunocytokine with PD-1 and IL2Rβγ binding domains, CTL immunophenotyping and phospho-proteomic analysis of intracellular metabolites showed significant upregulation of activation and functionality in CD8 T cells treated with RT + PD1-IL2v. Finally, we show the immunostimulatory response to RT + PD1-IL2v is significantly diminished with a concurrent lack of TCF+ CD8 T cell generation in the absence of functional IL15 signaling.

**Conclusions:** Our results are illustrative of a mechanism wherein unimpeded effector T cell activation through IL2Rβ signaling and Treg inhibition are necessary in mediating an anti-tumor immune response.

## Introduction

Despite treatment advances in surgery, chemotherapy, and radiation therapy (RT), pancreatic adenocarcinoma (PDAC) remains a devastating disease with a stagnant 5-year overall survival rate of 12%S. With few available treatment options, RT has been pursued as an adjuvant and neoadjuvant therapeutic in locally advanced and metastatic PDAC^1^. However, the widespread use of RT remains controversial, with radiation-induced fibrosis suggested as a possible correlate of worsened survival outcomes in some cases^2,3^. Further complicating treatment, many studies in the field of radiobiology have shown high dose RT delivered in large fractions increases the infiltration and activation of immunosuppressive regulatory T cells (Tregs) in locally advanced PDAC tumors^4^, and simultaneously fails to induce infiltration of CD8 T cells and natural killer (NK cells) critical for an anti-tumor immune response^4,5^, highlighting the narrow therapeutic utility for RT in the treatment of PDAC in current practice.

Despite these potential negative sequelae and the overall lack of immune infiltration following RT alone, studies have shown RT can enhance the effect of immunostimulatory treatments through the disruption of tumor vasculature and the release of proinflammatory cytokines and chemokines^6^, making it an attractive combinatorial option for immunomodulatory regimens. Moreover, recent studies have shown that the potential benefits of including RT in immunotherapeutic treatments extend beyond tumor lymphocyte infiltration. In our previous work, we demonstrated a synergistic effect of combination RT with the novel immunocytokine PD1-IL2v, which delivers variant interleukin-2 (IL-2) to PD1-expressing T cells while also inhibiting PD-1 signaling^7^. We established that the novel combination of RT + PD1-IL2v treatment not only increases cytotoxic T lymphocyte (CTL) frequency and polyfunctionality in both the tumor and draining lymph nodes, but also improves systemic immune surveillance and increases overall survival in orthotopic murine models of PDAC^7^. The RT + PD1-IL2v combination also resulted in the generation of a tumor-specific memory response that remained durable upon tumor rechallenge^7^, thus advocating for a role for RT in mediating a robust anti-tumor immune response when rational combinations are pursued.

Sharing many binding and signaling properties with PD1-IL2v, another emerging cytokine in the field of cancer immunobiology that has garnered much attention due to its ability to induce CD8 T cell memory and NK cell development and cytotoxicity is interleukin 15 (IL-15)^8^. Like PD1-IL2v and other IL2R-modulating agents, IL15 is a member of the common gamma chain family of cytokines and therefore has a strong impact on CTL functionality^8^. IL15 receptor signaling is unique in that it requires binding of IL15 to IL-15Rα on innate immune cells followed by subsequent cross-presentation of the IL15/ IL-15Rα complex to IL-2Rβγ on target cells^9^, allowing for tight regulation of endogenous IL15-mediated activation. To combat this signaling peculiarity, recombinant IL-15 complexed with IL-15Ra ex vivo is currently being explored as a therapeutic option due to its longer half-life, more profound expansion of CD8 T cells and NK cells, and overall reduction in tumor burden^10^. In this work, we evaluated the use of the IL-15/IL-15Ra complex (denoted IL-15c) in combination with RT while simultaneously managing RT-mediated Treg expansion using an aCD25 depletion antibody as a treatment strategy in PDAC tumors.

## Results

### Combination RT + IL15c is inefficacious; the addition of aCD25 to the RT + IL15c regimen prolongs overall survival and enhances CTL activation

To begin our analysis, we first sought to understand the translational potential of IL15 agonist therapies by interrogating RNA sequencing data from a previously published dataset of SBRT-treated PDAC patient tumor samples (GEO: GSE225767)^7^. We found that responders to SBRT alone exhibited elevated intratumoral levels of IL-15 transcripts prior to treatment (**Fig S1A**) and IL15 was among the most frequently appearing genes in the significantly upregulated pathways in responders relative to non-responders, appearing in 5 out of 10 pathways (**Fig S1B**).

Next, given the dependency of IL-15 signaling on the trans-presentation of IL-15Rα by innate cells to IL-2Rβγ on target cells (**Fig 1A**), we assessed the efficacy of our IL15 complex (see Methods) using an *in vitro* cytotoxicity assay (**Fig 1B**). We found that NKs stimulated with IL15c induced significantly more cancer cell death over control and that IL15c was more or equally effective in inducing cancer cell death compared to either IL2 or standard IL15 (**Fig 1C**). To characterize the effect of IL15c agonism *in vivo*, we next utilized Kras-driven orthotopic tumor PDAC models and treated tumor-bearing mice with RT and/or combination IL15c (**Fig 1D**). Given the central role of Tregs in mediating disease recurrence, particularly in the context of RT^5^, a treatment arm including a Treg-depleting aCD25 antibody was also included (**Fig 1D**). We found that neither RT + IL15c nor RT + aCD25 improved overall survival compared to RT alone (**Fig 1E**). However, the addition of aCD25 to RT + IL15c treatment did result in a significantly increased overall survival compared to RT (HR=3.8; p value=0.0017) and combination RT + IL15c (HR=4.9; p-value=0.0001), suggesting combination RT and IL15 agonism may have a clinical utility but only in the context of a Treg-deficient system (**Fig 1E**).

**Figure 1:**
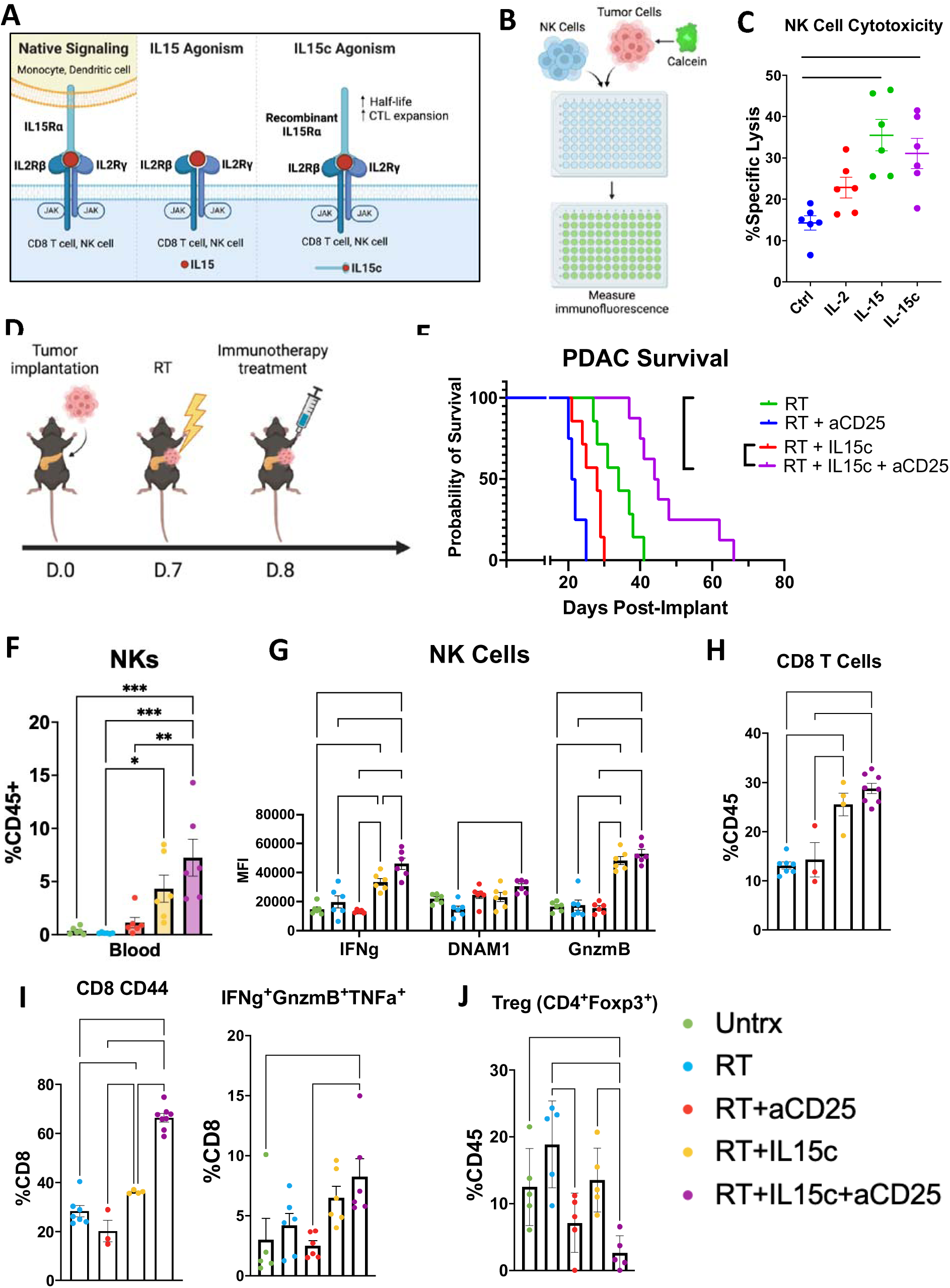
a. Schematic of IL15c binding properties b. Experimental design for NK cell cytotoxicity assay. n=6 per experimental group. c. In vitro percent specific killing of PK5L1940 cells by C57/Bl6 derived NK cells untreated and treated with IL-2, IL-15 and IL-15c as measured by calcein release. d. Experimental schematic of orthotopic survival study of Fig 1E. e. Kaplan-Meier survival analysis of orthotopically implanted pancreatic tumor-bearing C57/BL6 mice treated with RT (n=7) and IL15c (n=7), aCD25 (n=7), or IL15 + aCD25 (n=8). f. Flow cytometric analysis of NK cell frequency in the blood of C57/BL6 mice treated with RT and IL15c, aCD25, or IL15 + aCD25. g. Flow cytometric analysis of IFNg, DNAM1, and GnzmB MFI in peripheral NK cells harvested from C57BL/6 mice treated with RT and IL15c, aCD25, or IL15 + aCD25 h. Flow cytometric analysis of CD8 T cell frequency in the blood of C57/BL6 mice treated with RT and IL15c, aCD25, or IL15 + aCD25 i. Flow cytometric analysis of the frequency of CD44+ (left) and IFNg+/GnzmB+/TNFa+ (right) CD8 T cells in the blood of C57/BL6 mice treated with RT and IL15c, aCD25, or IL15 + aCD25. j. Frequency of intratumoral Tregs in C57/BL6 mice treated with RT and IL15c, aCD25, or IL15 + aCD25. n=5-6 per group for flow experiments Error bars represent SEM. ∗p < 0.05, ∗∗p < 0.01, ∗∗∗p < 0.001, ∗∗∗∗p < 0.0001.

As metastatic disease burden is highly prevalent in pancreatic cancer patients with over 50% of patients presenting with metastatic disease to the liver at the time of diagnosis^11,12^, we next explored the efficacy of the RT + IL15c + aCD25 regimen in a hemi-spleen metastatic model of PDAC using Kras-driven murine cell line^13^ (**Fig S1C**). As before, we observed a lack of response following RT + IL15c treatment relative to RT alone; however, the triple combination of RT + IL15c + aCD25 resulted in reduced metastatic disease burden and prolonged overall survival (HR=2.72; p-value=0.026) (**Fig S1D**).

To understand the mechanism of the improved response to combination RT + IL15c + aCD25 treatment, we performed flow cytometry on lymphocytes harvested from the blood and tumor of PDAC tumor-bearing mice. We found that animals treated with a regimen including IL15c had a significant expansion of circulating NK cells compared to control (**Fig 1F**), and NK cells treated with IL15c had significantly increased expression of IFNγ, DNAM-1, and GnzmB (**Fig 1G**). We also found that mice treated with RT + IL15c + aCD25 had significantly increased frequency of circulating CD8 T cells (**Fig 1H**), as well as increased CD44+ and IFNγ+/GnzmB+/TNFa+ polyfunctional CD8 T cells (**Fig 1I**). Importantly, mice treated with triple RT + IL15c + aCD25, the group with the best response rate (**Fig 1E**), had the lowest levels of intratumoral Tregs (**Fig 1J**), highlighting the importance of Treg depletion in mediating response to treatment.

### Response to RT + IL15c + aCD25 is dependent on CTLs and functional IL15 signaling

As sustaining the memory CD8 T cell population is a well-described defining feature of IL15 signaling^14^, we sought to determine whether the observed survival benefit following RT + IL15c + aCD25 triple combination treatment is dependent on the maintenance of the CD8 population. Utilizing the Rag1-/- mouse model lacking functional T and B cells (**Fig 2A**), we found that orthotopic tumor-bearing Rag1-/- mice treated with triple combination RT + IL15c + aCD25 had a median survival of 19.5 days as compared to 32 days in wildtype C57/BL6 mice treated with RT + IL15c + aCD25 (HR=4.09; p value=0.0001) (**Fig 2B**).

**Figure 2.**
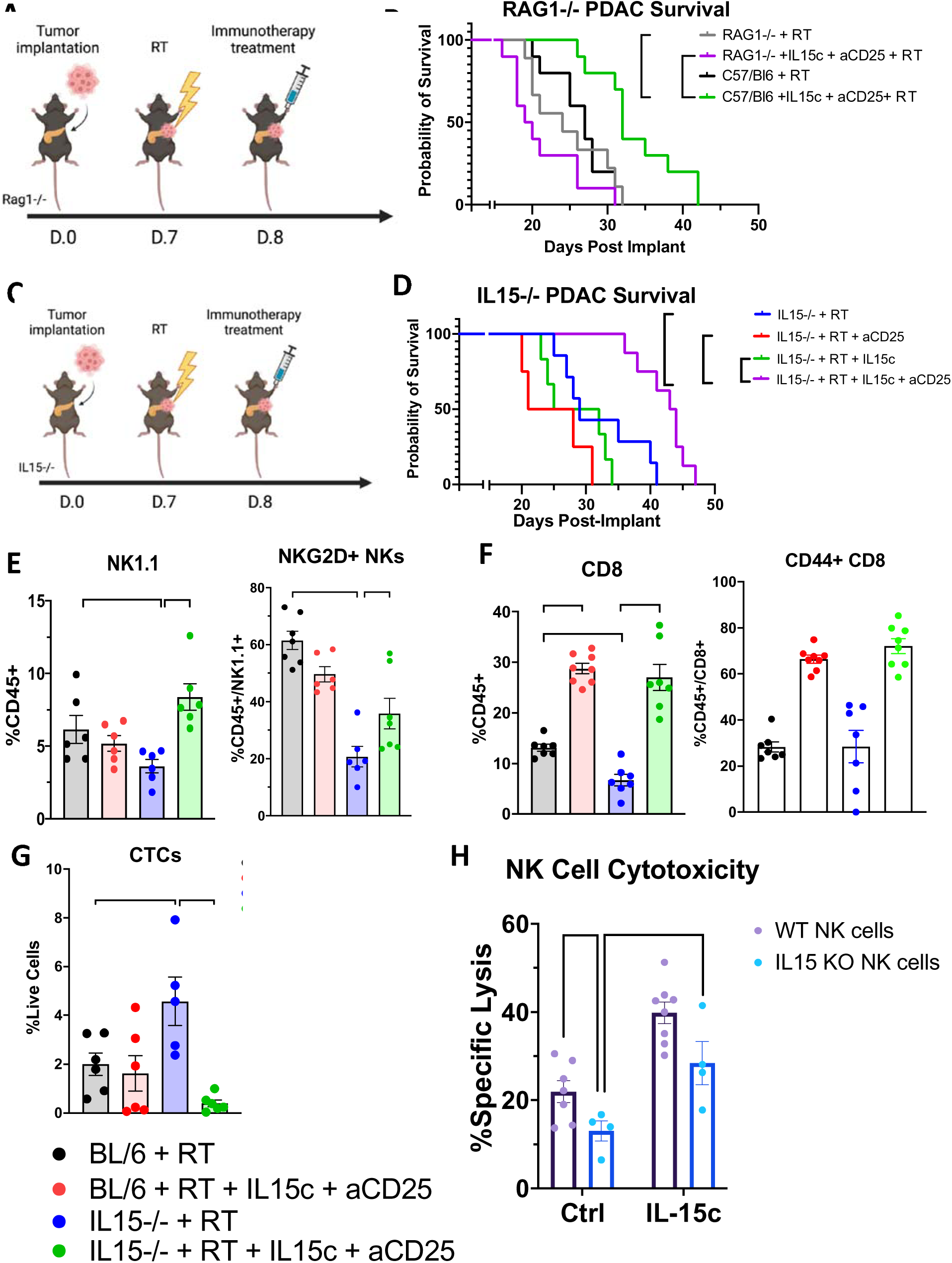
a. Experimental schematic of Rag1-/- survival study in Fig 2B b. Kaplan-Meier survival analysis of orthotopically implanted pancreatic tumor-bearing C57/BL6 mice treated with RT (n=10) and IL15c + aCD25 (n=10) and Rag-/- mice treated with RT (n=9) and IL15c + aCD25 (n=10). c. Experimental schematic of IL15-/- survival study in Fig 2D. d. Kaplan-Meier survival analysis of orthotopically implanted pancreatic tumor-bearing IL15-/- mice treated with RT (n=7) and IL15c (n=6), aCD25 (n=4), or IL15 + aCD25 (n=8). e. Flow cytometric analysis of NK cell frequency (left) and NKG2D+ NK cell frequency (right) in the blood of wildtype C57BL/6 and IL15-/- PDAC tumor-bearing mice treated with either RT alone or RT + aCD25 + IL15c. f. Flow cytometric analysis of peripheral CD8 T cell frequency (left) and CD44+ CD8 T cell frequency (right) in the blood of wildtype C57BL/6 and IL15-/- PDAC tumor-bearing mice treated with either RT alone or RT + aCD25 + IL15c. g. Flow cytometric analysis of circulating tumor cell (CTC) frequency in the blood of wildtype C57BL/6 and IL15-/- pancreatic tumor-bearing mice treated with either RT alone or RT + aCD25 + IL15c. h. In vitro percent specific killing of PDAC cancer cells by C57/BL6 and IL15-/- derived NK cells untreated and treated with saline control and IL15c as measured by calcein release. n=7-8 per group for flow experiments. Error bars represent SEM. ∗p < 0.05, ∗∗p < 0.01, ∗∗∗p < 0.001, ∗∗∗∗p < 0.0001.

We then sought to determine the dependence of the improved response to RT + IL15c + aCD25 treatment on IL15 signaling specifically. To do this, we utilized a globally deficient IL15 (IL15-/-) mouse model^15^ (**Fig 2C**). As seen in the wildtype system, RT + IL15c and RT + aCD25 resulted in no improvement in overall survival over RT alone. However, supplementation of aCD25 to the RT + IL15c treatment was able to significantly enhance survival benefit in an IL15-deficient system (HR=3.7; p-value=0.0031) (**Fig 2D**). Next, to characterize the circulating cellular populations contributing to the response to RT + IL15c + aCD25 in both WT C57/BL6 and IL15-/- mice, we used flow cytometric analysis of circulating immune cells and found a significant reduction in the frequency of NK and CD8 T cells in IL15-/- mice treated with RT compared to WT mice treated with RT (**Fig 2E-F**). However, both NK and CD8 T cell frequency and activation (as evidenced by NKG2D and CD44 expression, respectively) were restored with the addition of aCD25 and IL15c in the WT and IL15-/- mouse models (**Fig 2E-F**). Importantly, this restoration was associated with a concurrent decrease in CD45-/EpCAM+ circulating tumor cell (CTC) frequency, whereas the reduction in CTLs in the IL15-deficient system was associated with a significant increase in CTCs (**Fig 2G**). These data suggest that functional IL15 signaling is critical to enhancing CTL maturation and differentiation, reducing disease burden, and improving overall survival in response to RT + IL15c + aCD25 treatment.

Finally, to test the direct cytotoxic potential of CTLs following IL15c stimulation, we performed an *in vitro* cytotoxicity assay with NK cells isolated from WT and IL15-/- mice and treated them *in vitro* with IL-15c. We found that although NKs maturing in an IL15-deficient system have a lower baseline level of cytotoxicity, treatment with IL15c was able to significantly improve their cytotoxic potential (**Fig 2H**).

### RT + PD1-IL2v is superior to RT + IL15c + aCD25 in inducing immune activation, overall survival

The beta subunit of the IL2R receptor has recently garnered substantial interest as a target of selective agonism in anti-tumor immunity^9^. Our group and others have shown that PD1-IL2v, a novel immunocytokine consisting of an aPD-1 blocking antibody conjugated to an IL-2 variant optimized for IL-2Rβγ binding, robustly expands antigen-specific, TCF1+ anti-tumor T cells and leads to a significant survival benefit with RT and aPD-L1 combinations^7,16,17^. Because the IL15 signaling axis also utilizes IL2Rβ^9^, we next sought to characterize the differences in response to IL15c and PD1-IL2v treatments. Using an orthotopic model of PDAC (**Fig 3A**), we found that treatment with RT + PD1-IL2v resulted in a superior increase in overall survival relative to RT + IL15c treatment (HR=1.8, p-value=0.078) (**Fig 3B**).

**Figure 3.**
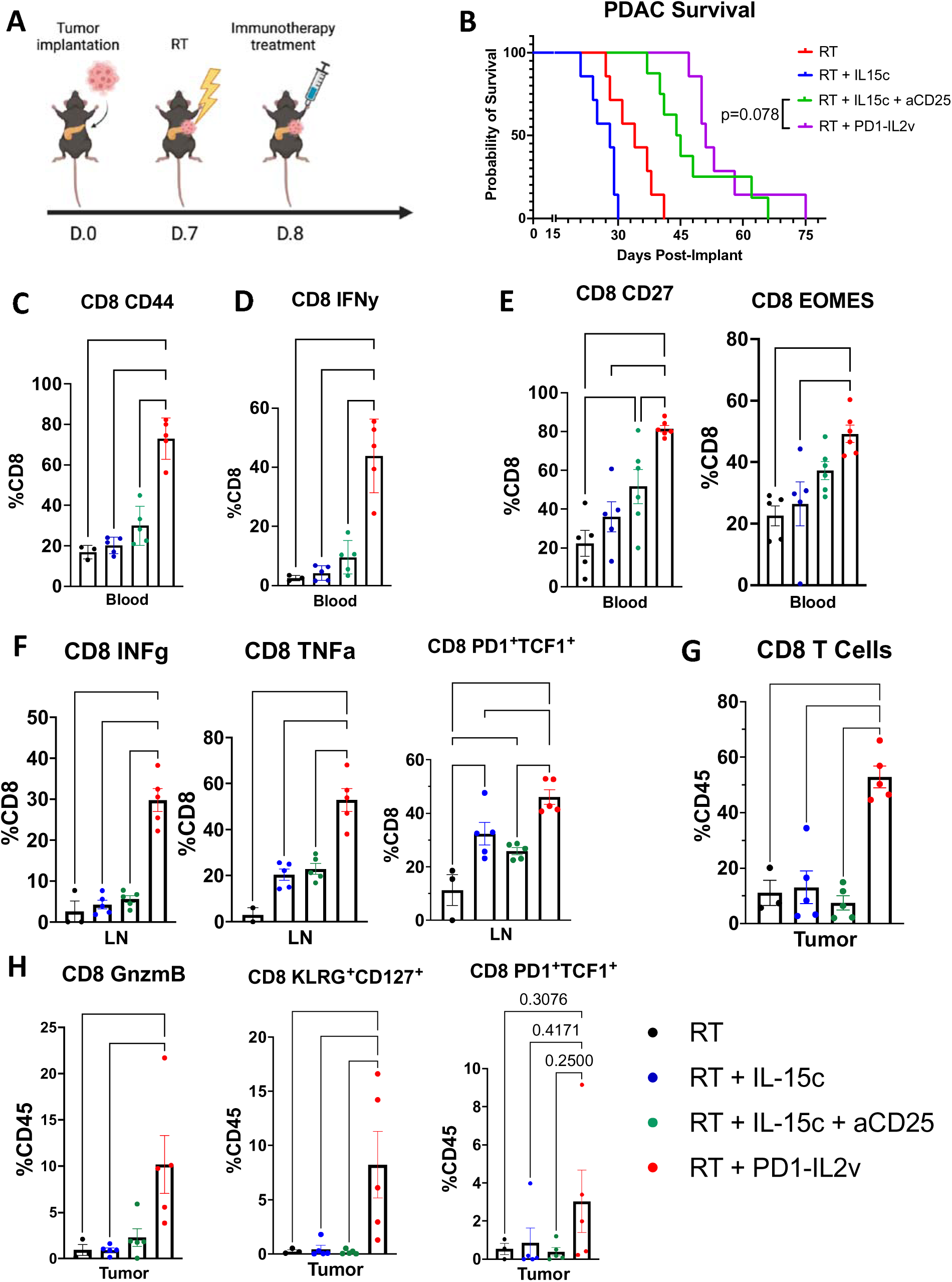
a. Experimental schematic of survival study in Fig 3B b. Kaplan-Meier survival analysis of orthotopically implanted PDAC tumor-bearing C57/BL6 mice treated with RT (n=7), RT + IL15c (n=7), RT + aCD25 + IL15c (n=8), and RT + PD1-IL2v (n=8). c. Frequency of circulating CD8+/CD44+ cells as a proportion of total CD8 T cells in orthotopic PDAC tumor-bearing mice treated with RT, RT + IL15c, RT + IL15c + aCD25, and RT + PD1-IL2v at 21 days post-implantation d. Frequency of circulating CD8+/IFNg+ cells as a proportion of total CD8 T cells in orthotopic PDAC tumor-bearing mice treated with RT, RT + IL15c, RT + IL15c + aCD25, and RT + PD1-IL2v at 21 days post-implantation e. Frequency of circulating CD8+/CD27+ (left) and CD8+/EOMES+ (right) cells as a proportion of total CD8 T cells in orthotopic PDAC tumor-bearing mice treated with RT, RT + IL15c, RT + IL15c + aCD25, and RT + PD1-IL2v at 21 days post-implantation f. Frequency of intramodal IFNg+ (left), TNFa+ (center), and PD1+/TCF1+ (right) CD8 T cells in the draining lymph nodes of orthotopic PDAC tumor-bearing mice treated with RT, RT + IL15c, RT + IL15c + aCD25, and RT + PD1-IL2v. g. Frequency of intratumoral CD8 T cells as proportion of total CD45+ cells in orthotopic PDAC tumor-bearing mice treated with RT, RT + IL15c, RT + IL15c + aCD25, and RT + PD1-IL2v. Frequency of intratumoral GnzmB+ (left), KLGR1+/CD127+ (center), and PD1+/TCF1+ (right) CD8 T cells in orthotopic PDAC tumor-bearing mice treated with RT, RT + IL15c, RT + IL15c + aCD25, and RT + PD1-IL2v. n=5-6 per group for flow experiments. Error bars represent SEM. ∗p < 0.05, ∗∗p < 0.01, ∗∗∗p < 0.001, ∗∗∗∗p < 0.0001.

Using multicompartmental flow cytometry, we then aimed to characterize the immune changes in response to each treatment. Through this analysis, we found that when compared to RT and RT + IL15c treatment, circulating CD8 T cells in animals treated with RT + PD1-IL2v more frequently expressed the activation markers CD44 (**Fig 3C**) and IFNγ (**Fig 3D**), and later in the disease course (24 days post-implantation), these animals also had a higher proportion of CD8 T cells expressing CD27 and EOMES (**Fig 3E**). Meanwhile, CD8 T cells in the draining lymph nodes of mice treated with RT + PD1-IL2v had significantly upregulated IFNγ and TNFα expression and an increased frequency in PD1+/TCF1+ stem-like memory CD8 T cells^18^ (**Fig 3F**). Within the tumor, RT + PD1-IL2v treated animals showed a profound increase in infiltrating CD8 T cells (**Fig 3G)**. Analysis of the activation and functional potential of tumor-infiltrating CD8s showed that those treated with RT + PD1-IL2v had significantly increased GnzmB+ and KLRG1+/CD127+ CD8 T cells, as well as a trend toward increased memory precursor effector CD8 T cells (MPECS)^19^ (**Fig 3H**), altogether suggesting that although RT + IL15c + aCD25 treatment increases the frequency of CTLs and depletes the immunoinhibitory Treg population, treatment with RT + PD1-IL2v is far superior in inducing CTL expansion and activation and increasing overall survival.

### RT + PD1-IL2v treatment induces significantly more phosphorylation of activation pathways in CD8 T cells than RT + IL15c + aCD25

Given the superior immunostimulatory effect of RT + PD1-IL2v relative to RT + IL15c + aCD25 despite operating on overlapping signaling axes, we hypothesized that while IL-15c and PD1-IL2v both interact with IL-2Rβγ on CD8 T cells, the differences in response to RT + PD1-IL2v and RT + IL15c treatment are due to disparate activation of CD8 T cell signaling cascades. To explore the phenotypic differences in CTLs following IL15c and PD1-IL2v administration, we performed phosphoproteomic analysis (see Methods, **Supp Fig 2A**) on circulating CD8 T cells harvested and isolated from tumor-bearing mice treated with RT, RT + IL15c + aCD25, and RT + PD1-IL2v (**Fig 4A**). We began our analysis with hierarchal and unsupervised clustering of samples from each treatment group which showed unique clustering patterns and distinct differences in phosphosite expression (**Fig 4 B-C**). Volcano plots of differentially expressed phosphosites were then generated and showed CD8 T cells treated with RT + PD1-IL2v had significantly more phosphorylated metabolites relative to control than those treated with RT + IL15c + aCD25, suggesting PD1-IL2v is superior to IL15c in initiating intracellular activation (**Fig 4D**).

**Figure 4.**
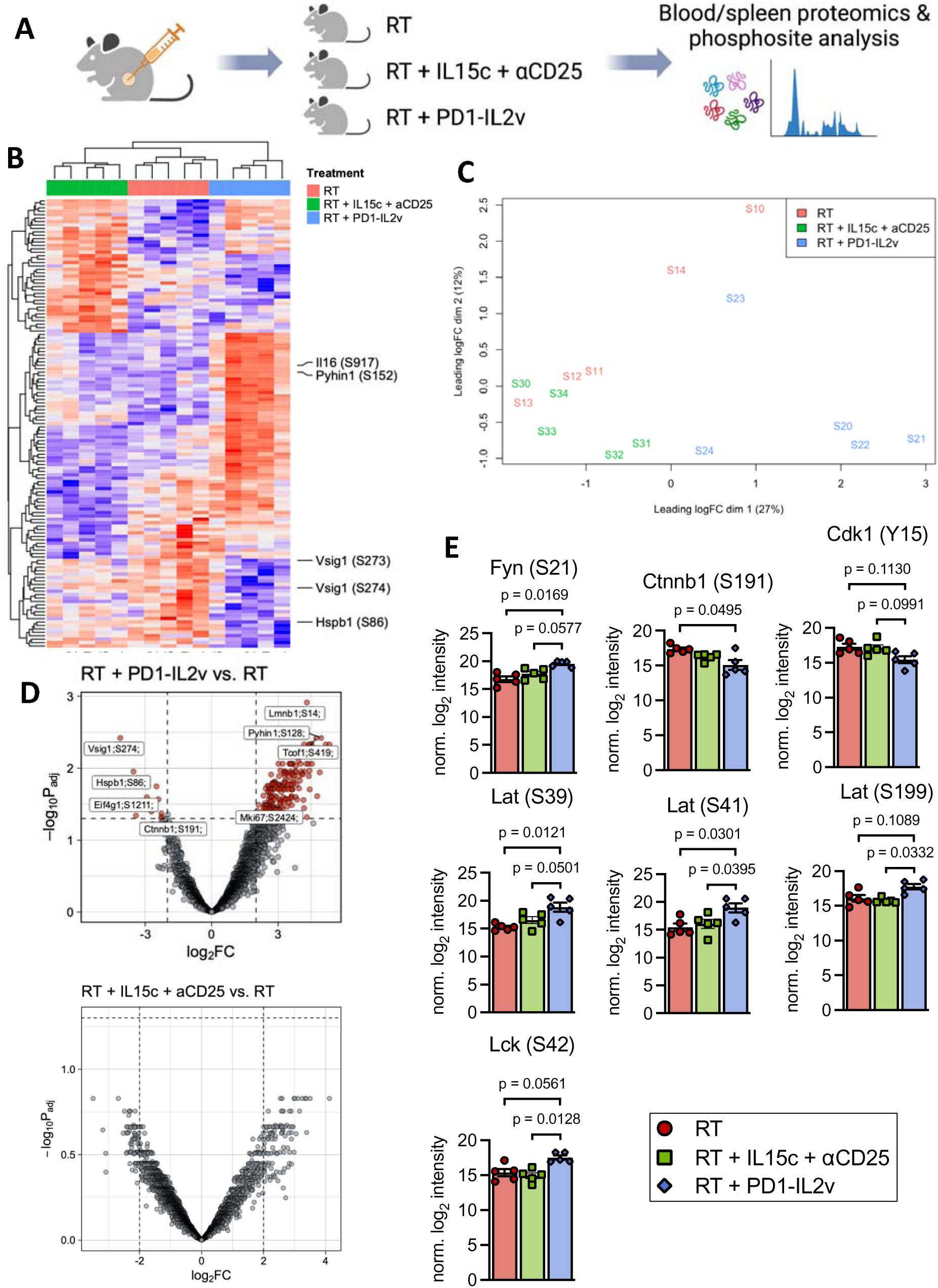
a. Experimental schematic of phosphoproteomic experiment b. Hierarchal clustering of phosphites from CD8s gathered from tumor-bearing mice treated with RT, RT + IL15c + aCD25, and RT + PD1-IL2v. c. PCA clustering analysis of phosphite samples from mice treated with RT, RT + IL15c + aCD25, and RT + PD1-IL2v d. Volcano plots of differentially expressed phosphites in CD8 T cells of mice treated with RT vs RT + PD1-IL2v (above) and RT vs RT + IL15c + aCD25 (below). Expression of phosphites related to T cell signaling and activation in CD8s harvested from tumor-bearing mice treated with RT, RT + IL15c + aCD25, and RT + PD1-IL2v. n=5 per group. Error bars represent SEM. ∗p < 0.05, ∗∗p < 0.01, ∗∗∗p < 0.001, ∗∗∗∗p < 0.0001.

We then shifted our analysis to individual phosphorylated proteins and found significant upregulation of a number of phosphosites in CD8 T cells from tumor-bearing mice treated with RT + PD1-IL2v, most notably those related to T cell signaling and activation pathways (**Figure 4E**). For example, we found increased phosphorylation at S42 of Lck, necessary for integrin-mediated signaling in T cells^20^, and increased S21 of Fyn, involved in cell migration^20^. Various residues also showed increased phosphorylation in Lat, downstream of Zap70, suggestive of superior TCR signaling (**Fig 4E**). Meanwhile, Ctnnb1 (beta-catenin) showed decreased phosphorylation at S191, which may indicate a polarization toward an effector rather than memory phenotype^21,22^. Phosphorylation was also decreased at the inhibitory residue Y15 of Cdk1, suggesting that T cell expansion may be increased and T cell anergy decreased in CD8 T cells treated with RT + PD1-IL2v^23,24^ (**Fig 4E**). Together, these data suggest that the unique stimulatory properties of the PD1-IL2v immunocytokine are highly effective at inducing the intracellular activation of CTLs and highlight the inherent limitations of endogenous signaling molecules as a form of anti-tumor treatment.

### Response to RT + IL15c treatment is not improved with the addition of immune checkpoint blockade

Given the divergent responses to RT + IL15c + aCD25 and RT + PD1-IL2v treatment, we next hypothesized that the inferiority of IL15-incorporating therapies is due to a lack of concurrent immune checkpoint inhibition. To test this claim, we orthotopically implanted pancreatic tumors into wildtype mice and treated them with RT + IL15c and RT + IL15c + aPD1. We then performed flow cytometry on peripheral and intratumoral immune populations and compared those findings to tumor-bearing mice treated with RT + PD1-IL2v (**Fig 5A**). We found RT + PD1-IL2v resulted in a trend toward reduced tumor size at day 23 post-implantation as compared to RT + IL15c + aPD1 (p-value=0.39) (**Fig 5B**).

**Figure 5:**
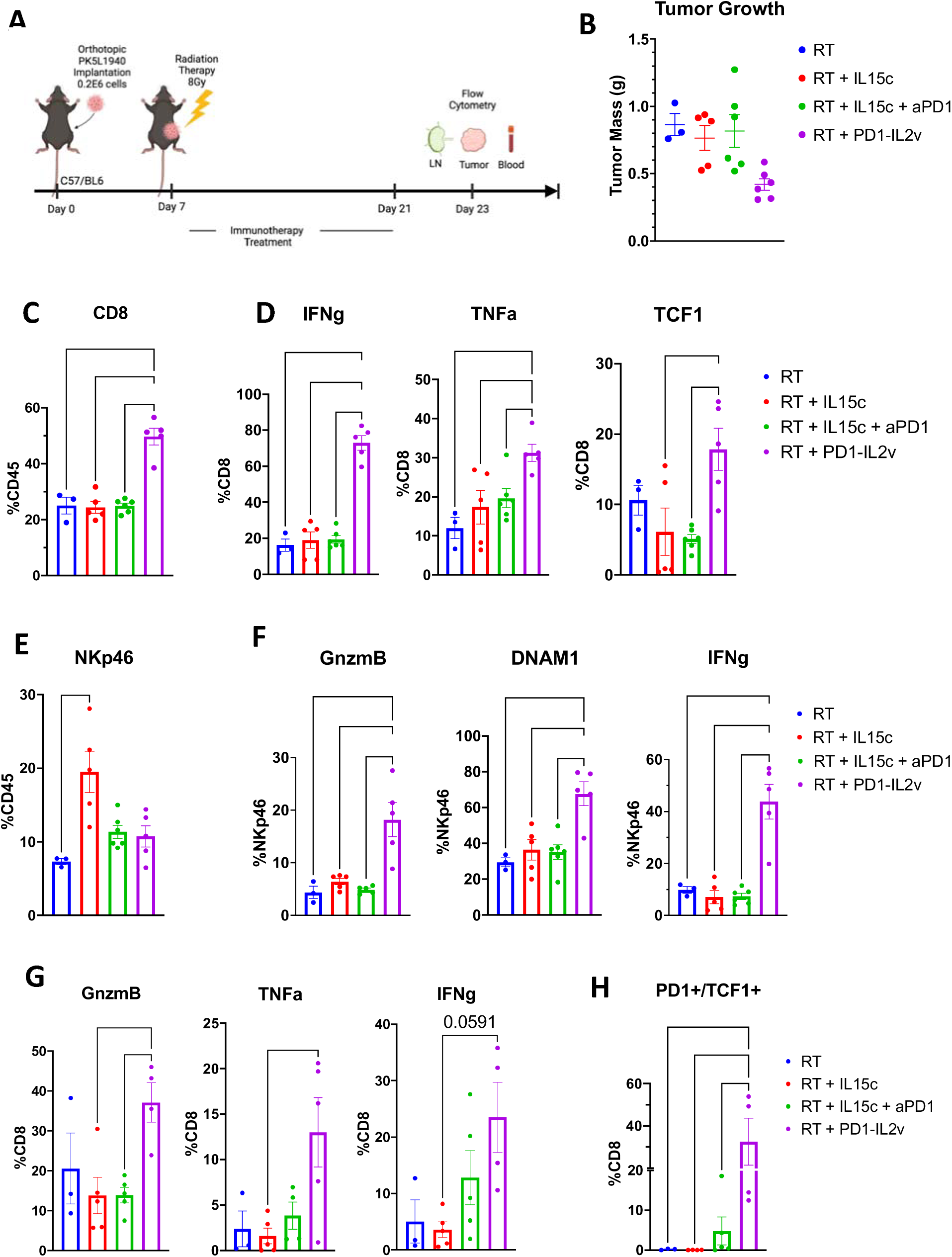
a. Experimental schematic of flow cytometric analysis of intratumoral and circulating immune cells. b. Mass of orthotopic pancreatic tumors treated with at day 23 post-implantation treated with combinations of RT, RT+IL15c, RT+IL15c+aPD1, and RT+PD1-IL2v. c. Frequency of CD8 T cells in the blood of orthotopically implanted PDAC tumor-bearing mice. d. IFNg (left), TNFa (center), and TCF1 (right) expression in circulating CD8 T cells of orthotopically implanted PDAC tumor-bearing mice. e. Frequency of NKp46+ NK cells in the blood of orthotopically implanted PDAC tumor-bearing mice. f. GnzmB (left), DNAM1 (center), and IFNg (right) expression in circulating NK cells of orthotopically implanted PDAC tumor-bearing mice. g. Frequency of GnzmB+ (left), TNFa+ (center), and IFNg+ (right) intratumoral CD8 T cells in orthotopically implanted PDAC tumor-bearing mice. Frequency of intratumoral PD1+/TCF1+ progenitor exhausted CD8 T cells in orthotopically implanted PDAC tumor-bearing mice. n=3-6 per group for flow experiments. Error bars represent SEM. ∗p < 0.05, ∗∗p < 0.01, ∗∗∗p < 0.001, ∗∗∗∗p < 0.0001.

Using flow cytometry to analyze peripheral immune populations, we found that mice treated with RT + PD1-IL2v had a significantly increased frequency of CD8 T cells relative to every other treatment group, including RT + IL15c + aPD1 (**Fig 5C**). Analysis of the activation state of these CTLs showed significantly more expression of IFNg, TNFa, and TCF1 (**Fig 5D**) in mice treated with RT + PD1-IL2v, suggesting enhanced activity and reduced exhaustion of CTLs in these animals. Of note, RT + PD1-IL2v resulted in similar levels of Treg frequency, as defined by CD4+/CD25+ cells, despite a significant increase in pro-inflammatory markers (**Fig S3A**). We also found that although RT + PD1-IL2v did not result in a significant expansion of the NK cell compartment as was observed in the RT + IL15c group (**Fig 5E**), RT + PD1-IL2v did induce significantly more expression of GnzmB, DNAM1, and IFNg in peripheral NK cells as compared to each other treatment arm (**Fig 5F**).

When looking at the intratumoral immune infiltrate, we found that the frequency of CD8 T cells was significantly increased in mice treated with RT + PD1-IL2v as compared to RT + IL15c + aPD1, as well as each other treatment group (**Fig S3B**). Moreover, tumor-infiltrating GnzmB+ and TNFa+ CD8 T cells were significantly increased and there was a trend toward increased IFNg production in mice treated with RT + PD1-IL2v (**Fig 5G**). Finally, mice treated with RT + PD1-IL2v had a significantly increased intratumoral frequency of PD1+/TCF1+ progenitor exhausted CD8 T cells, a CD8 subpopulation associated with increased stemness and self-renewal^25^ (**Fig 5H**). Altogether these findings suggest RT + PD1-IL2v combination treatment confers some functional advantage over RT + IL15c + aPD-1 despite shared PD-1 blockade and IL-2Rβγ signaling receptor agonism.

### IL15 signaling is necessary for durable response to RT + PD1-IL2v treatment

Because PD1-IL2v induces such a profound and systemic activation of CD8 T cells, one would reason that functional IL15 signaling is necessary to attain a maximal response to RT + PD1-IL2v treatment. As such, we next utilized our IL15-/- to test whether the response to RT + PD1-IL2v is dependent on IL15 signaling (**Fig 6A**). Using this model, we observed a significant reduction in overall survival following RT + PD1-IL2v treatment in IL15-/- mice when compared to their WT C57/BL6 counterparts (HR=4.47; p=0.0005) (**Fig 6B**). The addition of IL15c, however, rescued the effect of RT + PD1-IL2v treatment in the IL15-/- mouse model, as tumor-bearing mice treated with RT + PD1-IL2v + IL15c had a significantly improved survival over mice treated with RT + PD1-IL2v (HR=2.7; p-value=0.402) (**Fig 6C**). This suggests that PD1-IL2v requires an intact IL15 signaling axis to mediate a durable and systemic anti-tumor immune response.

**Figure 6:**
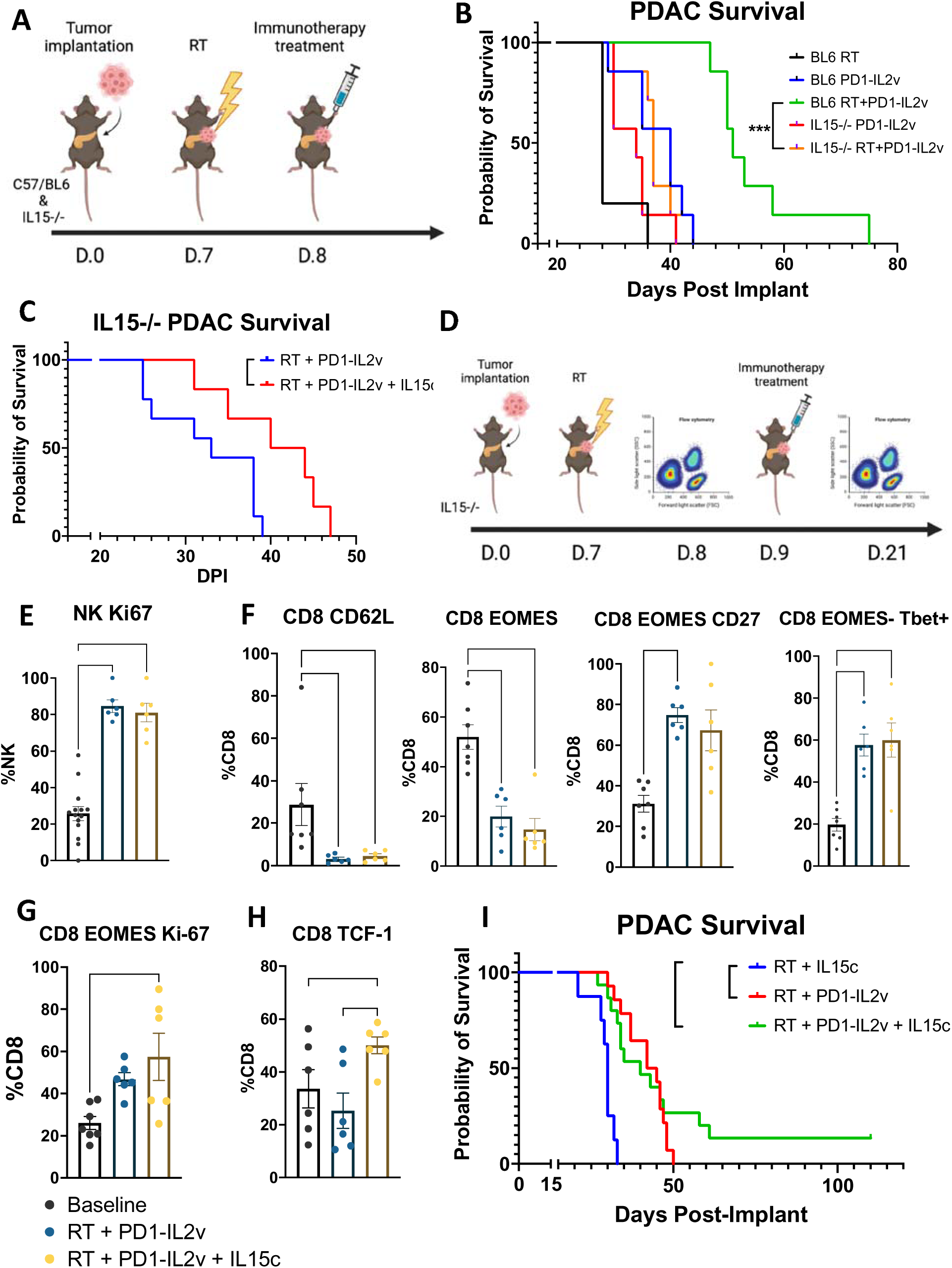
a. Experimental schematic of survival study in Fig 5B b. Kaplan-Meier survival analysis of orthotopically implanted PDAC tumor-bearing C57/BL6 and IL15-/- mice treated with PD1-IL2v or RT + PD1-IL2v (n=5-7 per group). c. Kaplan-Meier survival analysis of orthotopically implanted PDAC tumor-bearing IL15-/- mice treated with RT + PD1-IL2v (n=9) and RT + PD1-IL2v + IL15c (n=6). d. Experimental design of flow cytometry study in Fig 5E-H. e. Frequency of Ki67+ NK cells in the blood of IL15-/- orthotopic PDAC tumor-bearing mice treated with RT + PD1-IL2v (n=6) and RT + PD1-IL2v + IL15c (n=6) at baseline and following the initiation of treatment. f. Frequency of CD62L+, EOMES+, EOMES+/CD27+, and EOMES-/Tbet+ CD8 T cells in the blood of IL15-/- orthotopic PDAC tumor-bearing mice treated with RT + PD1-IL2v (n=6) and RT + PD1-IL2v + IL15c (n=6) at baseline and following the initiation of treatment. g. Frequency EOMES+/Ki67+ CD8 T cells in the blood of IL15-/- orthotopic PDAC tumor-bearing mice treated with RT + PD1-IL2v (n=6) and RT + PD1-IL2v + IL15c (n=6) at baseline and following the initiation of treatment. h. Frequency of TCF1+ CD8 T cells in the blood of IL15-/- orthotopic PDAC tumor-bearing mice treated with RT + PD1-IL2v (n=6) and RT + PD1-IL2v + IL15c (n=6) at baseline and following the initiation of treatment. Kaplan-Meier survival analysis of orthotopically implanted PDAC tumor-bearing C57/BL6 mice treated with RT + IL15c (n=8), RT + PD1-IL2v (n=14), and RT + PD1-IL2v + IL15c (n=16). Error bars represent SEM. ∗p < 0.05, ∗∗p < 0.01, ∗∗∗p < 0.001, ∗∗∗∗p < 0.0001.

To understand the impact of IL15 signaling on the response to RT + PD1-IL2v treatment, we then implanted orthotopic PDAC tumors into IL15-/- and used flow cytometry to characterize circulating immune populations before and after the initiation of RT + PD1-IL2v as compared to RT + IL15c + aCD25 treatment (**Fig 6D**). We found that mice treated with RT + PD1-IL2v RT + IL15c + aCD25 similarly induced expansion of Ki67+ NK cells (**Fig 6E**), decreased levels of CD62L+ and EOMES+ CD8 T cells, and increased levels of EOMES+/CD27+ and EOMES-/Tbet+ CD8 T cells (**Fig 6F**). Interestingly, however, only RT + PD1-IL2v + IL15c resulted in a significant increase in EOMES+/Ki67+ and TCF+ CD8 T cells relative to RT + PD1-ILv treatment (**Fig 6G-H)**, suggesting IL15 agonism may provide a unique contribution to the durable response via induction and maintenance of the central memory CD8 T cell population^26,27^.

Finally, we asked the question of whether RT + PD1-IL2v therapy could be further improved with the addition of concurrent IL15c administration. Although the addition of IL15c to RT + PD1-IL2v combination treatment did not significantly improve median survival over RT + IL15c, RT + PD1-IL2v + IL15c treatment did lead to a complete response in some animals, with 2 of 14 mice living past 110 days post-implantation free of tumor burden (**Fig 6I**).

## Discussion

Due to the overlapping intracellular involvement of JAK1, JAK3, and STAT3/5^9^, and importantly extracellular involvement of IL2Rβγ, many studies have been performed to understand the commonalities between IL2 and IL15 signaling, with results consistently showing that IL2 has a more prominent role in the initiation of an immune response functioning as a T cell growth factor and promoting the maintenance of self-tolerance, whereas IL15 is critical to a longer-lasting high-avidity T cell response involving the induction of CD8 memory T cells^9^. These effects would lead one to believe that cytokine-based therapies involving the administration of purified or recombinant IL2 and IL15 would elicit a significant anti-tumor effect. Such treatments, however, have been shown to come with their own set of challenges, including dose-limiting toxicity, off-target effects, and suboptimal efficacy^28^. To address these shortcomings, novel fusion proteins, such as IL15/IL15Ra complexes, have recently been developed and found to have enhanced biological activity, a longer half-life, and improved disease control in preclinical models^10^. In fact, IL15c therapies have shown so much promise that they are now making their way to the clinic, with the IL-15R agonist ALT-803/N-803 in combination with BCG now approved as a treatment modality in muscle-invasive BCG-unresponsive bladder cancer^29^, and IL15R agonist in combination with pembrolizumab or standard of care in stage III/IV NSCLC^30^. In this study, we sought to understand 1.) whether these novel approaches involving IL15 agonism can improve the response to RT in preclinical models of PDAC and 2.) whether IL15 agonism is superior to IL2 agonism, and in particular the novel PD1-IL2v immunocytokine^7^, in promoting an anti-tumor immune response and improving overall survival.

Our data show that unlike the combination of RT and PD1-IL2v, the addition of IL15c to RT is devoid of any additional antitumoral efficacy over RT alone and fails to confer a T effector immune response. Despite significantly enhancing NK and CD8 T cell activation, we show that the lack of effect following RT + IL15c treatment is driven primarily by regulatory T cells as further addition of a Treg-depleting aCD25 antibody to RT + IL15c elicited a significant anti-tumoral response approaching that of RT + PD-IL2v. Moreover, using the Rag1-/- mouse model lacking functional T cells, we show that Treg inhibition is necessary but not sufficient in mediating a systemic immune response to the RT + IL15c + aCD25 treatment regimen and response requires the presence of activated CTLs. Translationally, these findings advocate for the concurrent targeting of Tregs in therapies incorporating IL15 agonism as the beneficial effects of IL15 agonism are significantly impeded by Treg-mediated immunosuppression.

Although the topological structures of IL2Rα (CD25) and IL15Rα (CD215) have been shown to be nearly identical^9^, the ability of PD1-IL2v to preferentially stimulate IL2Rβγ on CTLs while subverting IL2Rα agonism on Tregs is one possible explanation for its superior efficacy^7^. In this work, through mechanistic studies characterizing the differences in response to IL15c and PD1-IL2v, we also identified a key role for CD8 stem cells, without which a durable immune response is not possible and without which the immunocytokine PD1-IL2v cannot generate an anti-tumoral systemic immune response. Interestingly, by utilizing an IL15-/- mouse model, we show that this potent CD8-driven antitumoral immune response is dependent on the presence of IL15 and, without it, no TCF+ CD8 T cells can be generated and all response to RT + PD1-IL2v combination treatment is lost. Moreover, our phospho-proteomics analysis of CD8 T cells derived from RT + PD1-IL2v and RT + IL15c + aCD25 tumor-bearing mice, in concordance with findings by others, show that although the IL2 and IL15 signaling pathways share the IL2Rβγ complex, treatment of T cells with high concentration IL2 and IL15 results in differential intracellular signaling and tnf, Ifng, and Il2ra gene expression^31^. Together, these results identify the generation of CD8 stem cells as a necessary pre-requisite for the generation of effective anti-tumoral immune response and are suggestive of a mechanism wherein unimpeded effector T cell activation following IL2 agonism is necessary in the acute anti-tumor immune response, whereas IL15 signaling plays an important role in the generation of a systemic and durable immunity.

Of note, the idea of utilizing a bifunctional signaling molecule to provide IL15-mediated immunostimulation while simultaneously inhibiting immunosuppression does have precedent in the literature. In a recent study by Liu et. al., the group describes the creation and biological activity of a heterodimeric bifunctional fusion molecule composed of soluble TGFβ-R and IL15/IL15Rα domains (HCW9218) as a novel immunotherapeutic agent with both IL15-mediated immunostimulatory and TGFb antagonistic activities^32^. They show that this molecule significantly enhances NK and CD8 T cell activation and infiltration, reduces circulating plasma TGFβ levels, and improves disease control. Meanwhile, Xu et. al. recently generated an IL15-mutein and PD1-specific antibody fusion complex (aPD1-IL15m) and demonstrated that this molecule was more biologically active than either of its components alone. They found that aPD1-IL15m robustly enhances the proliferation, activation, and cytotoxicity of CTILs and improves anti-tumor immunity^33^. The results described in our work would seem to validate these studies while going further to establish a role for radiation in such combinatorial approaches and emphasizing the importance of IL2-mediated generation of TCF1+ CD8 T cells in generating a durable response to treatment.

Collectively, our data reinforce the now widely understood differences between IL2 and IL15 signaling and highlight the important differences between IL2 and IL15 in mediating the acute and chronic anti-tumor immune response. These results also advocate for the use of RT when used in the appropriate setting and will help guide future clinical trials exploring IL2Rβ-modulating agents in novel combinations.

## Acknowledgements

Sana D. Karam is funded by the following grants: R01DE028529, R01CA28465, R01DE028529, 1P50CA261605-01, V Foundation. Jacob Gadwa is funded by the following grant: F31 DE033887-01 This work was also supported by the Wings of Hope Foundation for Pancreatic Cancer Research.

## Materials and Methods

### Patients and samples

Written consent was obtained for all tumor sample collection. Studies were performed in accordance with U.S. Common Rule and approved by institutional review board. Patient archival tumor samples were identified and obtained from the University of Colorado PSR biorepository and collected per COMIRB13-0315. Samples were selected from all borderline resectable pancreatic cancer patients seen through University of Colorado pancreatic multidisciplinary clinic between 1/2013–12/2018 and were treated with FOLFIRINOX or gemcitabine based neoadjuvant chemotherapy and stereotactic body radiotherapy (SBRT). Following neoadjuvant therapy, all patients received surgery followed by further adjuvant chemotherapy.

### Cell lines and reagents

PK5L1940 mouse pancreatic adenocarcinoma cell line was kindly provided by Dr. Michael Gough (Providence Cancer Institute, Portland, OR). FC1242 mouse pancreatic adenocarcinoma cell line was kindly provided by Dr. David Tuveson (Cold Spring Harbor Laboratory, Cold Spring Harbor, NY). Cell lines were passaged in RPMI1640 supplemented with 10% FBS every 2–3 days at a density of 1:4-1:10. Cells were not allowed to grow beyond passage 30.

### Mice

Female C57BL6 (6-8 weeks old) and Rag1-/- (B6.129S7-Rag1tm1Mom/J) (6-8 weeks old) were purchased from Jackson Laboratories (Indianapolis, In, USA). IL15-/- mice were kindly provided by Dr. Ross Kedl (University of Colorado, Denver, CO).

All mice were cared for in accordance with the ethical guidelines and conditions set and overseen by the University of Colorado, Anschutz Medical Campus Animal Care and Use Committee. Protocols used for animal studies were reviewed and approved by the Institutional Animal Care and Use Committee at the University of Colorado, Anschutz Medical Campus.

Experimental unit size was determined using historical results to maximize statistical significance and minimize animal death (n=3-6 per group for flow cytometry studies; n=7-12 for survival studies). Mice were randomized into treatment groups on day 6 post-implantation, prior to the initiation of treatment. All experimental mice were included in survival and immunophenotyping studies.

### Local and metastatic pancreatic adenocarcinoma implantations

Local orthotopic implantations were conducted by first anesthetizing mice using isoflurane and making a 1 cm incision in the left subcostal region. Mouse pancreata were located, externalized, and injected with 200,000 PK5L1940 or FC1242 KPC cells suspended 1:1 in Matrigel (Corning, Corning, NY). Pancreata were then reintroduced into abdomen and mice peritoneum and skin were closed. Protocol described in further detail^34^ . Survival and flow cytometric in vivo studies were conducted and analyzed separately.

Metastatic orthotopic implantations were conducted as above with spleen externalization following subcostal incision. Spleens were first ligated with horizon clips and 1 hemispleen was injected with 200,000 FC1242 cells suspended in 50μl 10% RPMI followed by washout injection of 50 μl PBS. Pancreatic vessels were then ligated with horizon clips and hemispleen was excised prior to closure of peritoneum and skin. Metastatic implantation described in further detail^13^. For cancer specific mortality, mice determined to have died from other causes were excluded from the analysis.

### *In vivo* drug administration

aCD25 (Roche Pharmaceuticals, 3mg/kg), PD1-IL2v (muPD1-IL2v, Roche Pharmaceuticals, 0.5mg/kg)^16^, and DP47-IL2v (muDP47-IL-2v) (0.5mg/kg) were administered intraperitoneally once per week beginning one week after tumor implantation. aPD-1 (BioExcell, 10 mg/kg) was administered intraperitoneally twice per week beginning one week after tumor implantation consistent with the previously published protocol^7^. IL15c was generated by mixing recombinant IL-15 with recombinant IL-15Ra at a m/m ratio of 1:10 and incubated for 30 minutes at 37C prior to administration. IL-15c (0.4mg/kg) was administered intraperitoneally twice per week beginning one week after tumor implantation.

### Flow cytometry

For flow cytometric analysis of tumor tissue, tumors were digested into single-cell suspension as previously reported^35^. Briefly, tumors were finely cut and incubated in HBSS solution with Collagenase III (Worthington) at 37°C. After incubation, tumors were passed through a 70 μm nylon mesh. The resulting cell suspension was centrifuged and re-suspended in red blood cell (RBC) lysis buffer for 5 minutes. RBC lysis buffer was deactivated, cell suspensions were centrifuged, re-suspended, and counted using an automated cell counter. Tumor-draining inguinal lymph nodes and spleens were processed into single-cell suspensions as above. For flow cytometric analysis, cells were plated in 24-well plates and cultured for 4 hours in the presence of monensin, PMA, and ionomycin to stimulate cytokine production and block Golgi transport. Cells were then blocked with anti-CD16/32 antibody. Where necessary, cells were fixed and permeabilized prior to staining using the FoxP3 Fixation/Permeabilization protocol (eBioscience). Samples were run on the Cytek Aurora Spectral Cytometer at the Barbara Davis Center at the University of Colorado Diabetes Research Center (NIDDK grant #P30-DK116073). Data were analyzed using FlowJo Analysis software. Populations were visualized using FlowSOM within Cytobank software.

### Radiotherapy

Image-guided radiotherapy was performed using the X-Rad SmART small animal irradiator (Precision X-Ray, North Bradford CT) at 225kVp, 20mA with 0.3 mm Cu filter. Mice were positioned in the prone orientation and a CT scan was acquired. Radiation was delivered at a dose rate of 5.6Gy/min. A single 8 Gy dose of X-ray radiation was delivered to mouse pancreata using 10mm square beam with field edges at mouse midline and below left ribs. Monte-Carlo simulation was performed using SmART-ATP software (SmART Scientific Solutions, Maastricht, Netherlands) with a CBCT scan of one mouse to determine the appropriate time and current. All mice received identical treatment after repositioning by fluoroscopy. For all in vivo experiments, radiation was given at 7 days post-implantation as previously described^7^.

### In Vitro Calcein Release Assay

Cytotoxicity assay was performed by pre-staining PK5L1940 pancreatic cancer cell lines using a 2mg/ml calcein solution and isolating NK cells according to the manufacturer’s protocol. Briefly, cancer cells were incubated in calcein-containing media for 30 minutes at 37C and subsequently plated in a 96-well plate at a 2:1 NK cell to cancer cell ratio. The reaction mixture was then allowed to incubate at 37C for 4 hours. Following the incubation period, supernatant was collected and cancer cell release of calcein was quantified by fluorescence (Ex: 485nm/EM: 530 nm) Protocol described in further detail^36^.

### Phospho-proteomics/phospho-array

Samples were prepared for mass spectrometry using S-Trap^TM^ micro filters (Protifi, Huntington, NY) according to the manufacturer’s protocol. Phosphopeptides were enriched using the Fe-NTA Phosphopeptide Enrichment Kit (ThermoFisher Scientific) and subjected to liquid chromatography tandem mass spectrometry using a NanoElute liquid chromatography system coupled with a timsTOF SCP mass spectrometer (Bruker, Germany). Raw spectra were interpreted against the UniProt *Mus musculus* protein sequence database using MSFragger with a false discovery rate of <1.0%. Phosphoproteomics data was analyzed using R as follows. Intensity data collected with phosphosites in rows and samples in columns was filtered to only include rows with phosphorylation events: rows with assigned modifications including a weight change +79.9663 Da. Some data rows included multiple modifications per quantified peptide; these rows were separated vertically so that each row would include intensity of a single peptide phosphorylation event per row. Positions of modifications were shown in original data relative to the start of each peptide sequence identified; these positions were converted to an absolute position relative to the entire peptide sequence of matched protein ID, which allowed for identification with known phosphorylation events documented in the literature. Rows at this point that mapped to the same protein-phosphosite pair were combined by sum, such that each row showed sample intensities for sites of the format *<*GENE symbol*>; <*RESIDUE*> <*POSITION*>;* for phosphorylation events at serine, threonine, or tyrosine residues. Further data shaping as follows: Rows were filtered out if in intensity was constant across the entire row for all samples. Values with expression zero were interpreted as missing not at random, as missing values occurred with lower intensity, suggesting dropouts for lack of detection under a low threshold (see density plot in **Fig S2A**). Rows were then only kept if non-missing intensity was detected in at least 50% of replicates (at least 3 of 5 replicates in a group) within at least one condition (RT, RT + IL15c + aCD25, or RT + PD1-IL2v) using the selectGrps() function with R package PhosR^37,38^. Variance-stabilizing normalization was applied with values returned on the log2 scale with R package vsn^39^ with functions vsn::vsnMatrix() and vsn::predict() with log2scale = TRUE. Missing values were imputed conservatively, with assumption of left-censored distributions in each sample, using R package imputeLCMD with function impute.MinProb() with q = 0.01 and tune.sigma = 1 to draw minimum values. BH-adjusted p-values were generated with R package limma using lmFit() and eBayes().

## Statistical significance

Quantitative analyses were performed using a two-sided Student’s t-test, Mann-Whitney test, One-Way ANOVA with multiple comparisons, or the Mantel-Cox test for survival using GraphPad Prism. p-values of <0.05 were considered statistically significant. Statistical outliers were identified by ROUT method and removed prior to analysis. All experimental conductors were aware of treatment group allocations.

## Author Contributions

All authors contributed to the study conception and design. Material preparation, data collection, and analysis were performed by MP, JG, CH, and MK. The first draft of the manuscript was written by MP and all authors commented on previous versions of the manuscript. All authors read and approved the final manuscript.

## Data Availability

Genomic data, including human PDAC RNA sequencing and phosphoproteomics, as well as supporting materials including R code used to analyze phosphoproteomic data, are available from the corresponding author upon reasonable request.

