## Supplemental Data for "IL15/IL15Rα complex induces an anti-tumor immune response following radiation therapy only in the absence of Tregs and fails to induce expansion of progenitor TCF1+ CD8 T cells"

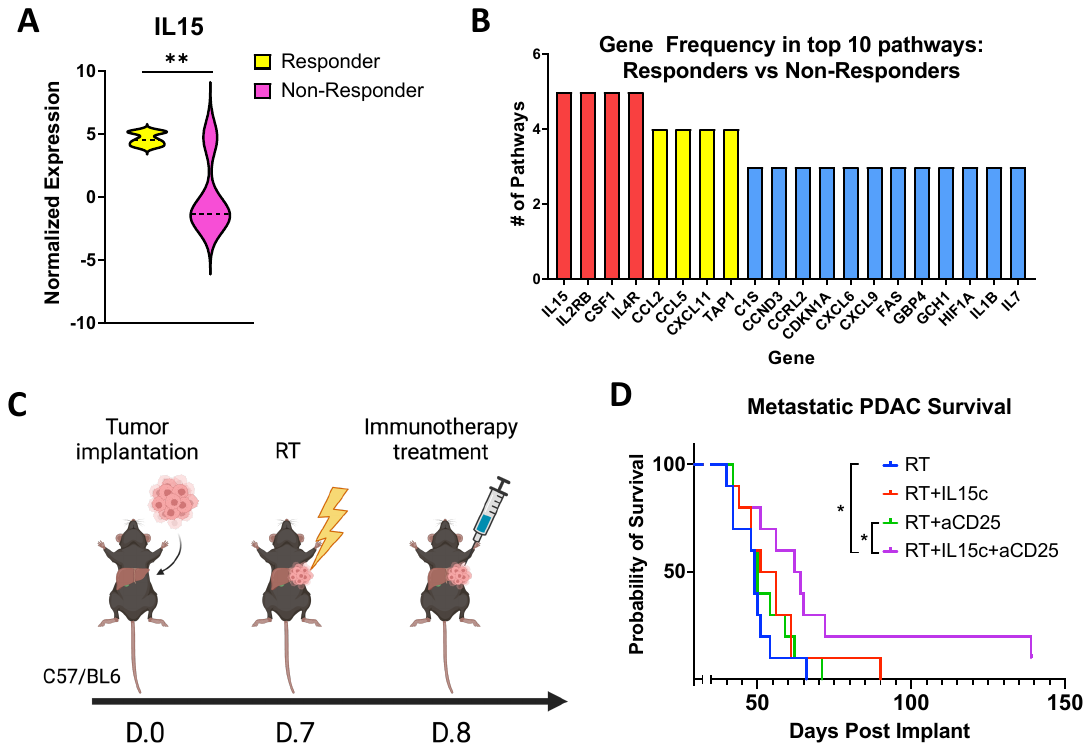

Supplementary Figure 1

Supplementary Figure 1:

a. Normalized expression of IL15 mRNA in responders and non-responders to RT treatment in human PDAC samples.
b. Genes appearing in the most frequently upregulated pathways in responders relative to non-responders to RT treatment in human PDAC samples.
c. Experimental schematic of metastatic orthotopic PDAC survival studies.
a. Kaplan-Meier survival analysis of tumor-bearing mice treated with RT and IL15c, aCD25, or IL15 + aCD25 using a hemispleen metastatic model of PDAC. n=10 per group. ∗p < 0.05, ∗∗p < 0.01, ∗∗∗p < 0.001, ∗∗∗∗p < 0.0001.

Supplementary Figure 2

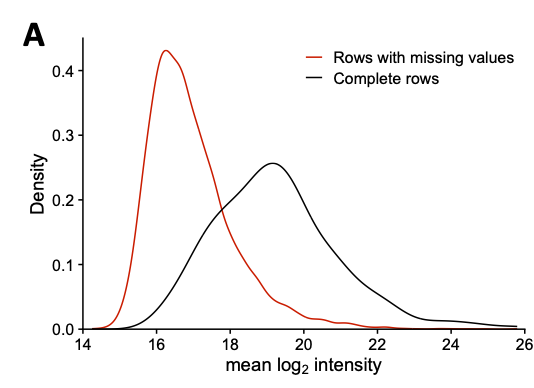

Supplementary Figure 2:

a. Density plot of phosphosite intensity for phosphosites split by presence or absence of missing values for some replicates, demonstrating a bias toward lower intensity found in phosphosites with missing values.

Supplementary Figure 3

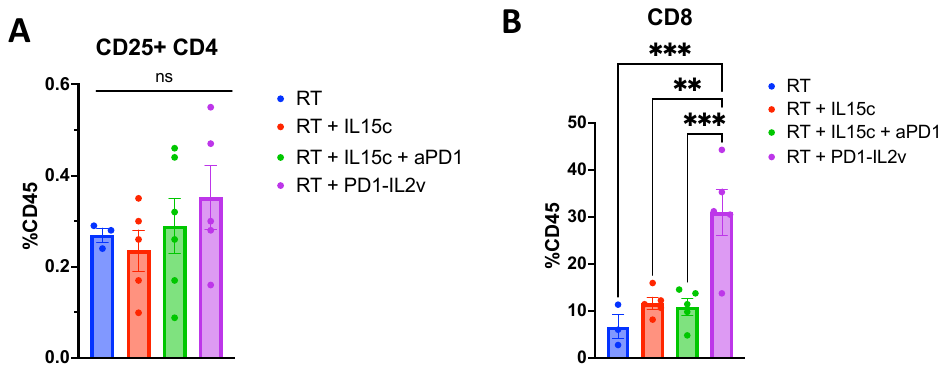

Supplementary Figure 3:

a. Frequency of circulating CD25+ CD4 T cells in the blood of orthotopically implanted PDAC tumor-bearing mice treated with combinations of RT, RT+IL15c, RT+IL15c+aPD1, and RT+PD1-IL2v.
b. Frequency of intratumoral CD8 T cells in orthotopically implanted PDAC tumors treated with combinations of RT, RT+IL15c, RT+IL15c+aPD1, and RT+PD1-IL2v. n=5-6 per group. ∗p < 0.05, ∗∗p < 0.01, ∗∗∗p < 0.001, ∗∗∗∗p < 0.0001.
